# The Pollen Tube Penetrates the Synergid Cell by Formation of a Peritubular Membrane

**DOI:** 10.1101/2024.07.10.602759

**Authors:** Nicholas Desnoyer, Marta Belloli, Stefano Bencivenga, Philipp Denninger, Ueli Grossniklaus

## Abstract

In flowering plants, successful reproduction relies on an exchange of signals between synergids and pollen tubes (PTs), mediating the invasion of a synergid by the PT, which then ruptures and releases two sperm cells to effect double fertilization. However, how exactly the PT invades the receptive synergid is unknown as the spatial relationship between these two cells is unclear. To better understand this process we performed 3D live imaging of PT reception in *Arabidopsis thaliana*. Upon arrival at the filiform apparatus (FA), a region rich in membrane folds at the micropylar pole of the synergids, the PT gradually deforms the FA before it rapidly grows into the receptive synergid. Upon penetration, the membrane of the receptive synergid invaginates and envelopes the PT. We termed this newly discovered structure the peri-tubular membrane (PRM). We show that, in *feronia* mutants disrupting PT reception, the PT still enters the receptive synergid, forming a normal PRM. This results in extensive invagination of the synergid membrane without sperm release. We show that PRM formation is associated with a cytosolic calcium ([Ca^2+^]_cyt_) spike of high amplitude in the PT and flooding of [Ca^2+^]_cyt_ in the synergids. In PTs lacking AUTOINHIBITED Ca^2+^ ATPASE9 activity, PTs have lower amplitude [Ca^2+^]_cyt_ spiking and the PTs frequently fail to penetrate the synergid. Our findings suggest that synergid penetration and the non-cell autonomous control of PT rupture are distinct regulated processes required for fertilization in flowering plants.

## Introduction

During flowering plant reproduction, the pollen grain, or male gametophyte, germinates a pollen tube (PT) that invasively grows through the tissues of the pistil to finally deliver its two sperm cells to the female gametes, which are harbored within the female gametophyte (1). In *Arabidopsis*, the germinated PT first penetrates the cell wall of a papillae cell of the stigma, where the cytoskeletal dynamics and mechanical properties of the papillae impact PT growth direction (2). PT penetration of the papillae represents an interspecific reproductive barrier and is partly regulated by FERONIA (FER) (3, 4), a member of the *Catharanthus roseus* RECEPTOR-LIKE KINASE1-LIKE family, that perceives secreted peptide ligands of the RAPID ALKILINIZATION FACTOR (RALF) family (5–9). Following penetration of the papillae, the PTs grow through the transmitting tissue of the style and septum into the gynoecium (10). Here, FER/RALF signaling along with ovular signals regulate the emergence of a single PT from the transmitting tract onto the epidermal layer of the septum into the locules of the gynoecium (11, 12). The PT is then guided by female gametophytic signals along the funiculus of the ovule into the micropyle, where it comes into contact with the filiform apparatus (FA) of the synergids, a region of dense membrane invaginations and cell wall thickenings (13–15). Here, the synergid-expressed FER receptor kinase coordinates intercellular communication between the PT and the synergid in a process called PT reception. Upon arrival at the FA, the PT grows very slowly for about 30 minutes (phase I), before it rapidly grows either into or around the receptive synergid (phase II) (3, 16, 17), leading to PT rupture, release of the two sperm cells, and death of the receptive synergid (18–20). PTs encountering a *fer* mutant synergid will continue to grow without rupture, resulting in extensive PT overgrowth and coiling inside or around the female gametophyte (3, 4).

In addition to regulating invasive growth of the PT through the female reproductive tissues, *FER* also plays a role as a susceptibility factor for powdery mildew, as *fer* mutant epidermal cells are poorly penetrated by fungal hyphae (21). Furthermore, NORTIA (NTA), a membrane protein of the MILDEW RESISTANCE LOCUS O (MLO) family, acts downstream of FER in synergids and its polar deposition into the FA is required for proper PT reception (21–23). In the context of powdery mildew infection, polar localization of MLO proteins is important to prevent entry of the hyphae into the host cells (24). In contrast to pathogenic fungi, the invasive growth of endophytic fungi, such as arbuscular mycorrhizae, is assisted by the host plant (25). Contact of mycorrhizal fungi with root epidermal cells triggers nuclear calcium oscillations, and the host cells form an accommodating structure, the pre-penetration apparatus, which directs hyphal growth through the root (26–28). As mycorrhizal hyphae penetrate epidermal and cortical cells to form arbuscules, they are surrounded by the plant’s peri-arbuscular membrane that facilitates nutrient exchange between the symbionts. In *Medicago truncatula* and *Lotus japonicus*, several mutants have been identified that are deficient in the formation of the pre-penetration apparatus and fungal entry, demonstrating that this is a host-regulated process and that the pre-penetration apparatus is essential for arbuscule formation (26, 29–31). While *Arabidopsis* does not form a symbiotic association with mycorrhizal fungi, the invasive growth of PTs, which is regulated by FER that also controls fungal invasion, begs the question of whether FER signaling regulates penetration of the receptive synergid by the PT and whether any accommodating structures similar to the to pre-penetration apparatus and peri-arbuscular membrane are formed during PT reception.

It is currently unclear where exactly the PT grows prior to rupture and whether PT coiling in *fer* synergids is a consequence of the PT failing to penetrate and enter the receptive synergid or of failing to trigger rupture once it is within the receptive synergid. Aniline blue staining of PTs coiling in *fer* mutants showed that PTs often form spiral structures that may be outside the female gametophyte (Fig. S1). A previous study in the wild type indicated that the PT grows around the receptive synergid and penetrates it near the synergid hooks, a site distinct from the filiform apparatus (16). Other investigations on fixed tissue suggested that the PT enters the receptive synergid via the FA (3, 17), while in live imaging of PT reception in *fer* mutants, it was unclear if the PTs coiled around or inside the synergids (18). Clearly, a better understanding of the events during PT reception requires resolving where the PT enters the receptive synergid and how this relates to rapid PT growth, PT rupture, and death of the receptive synergid. But so far, no high resolution imaging of the penetration process has been performed, although more than ten studies implemented live imaging of PT guidance, PT reception, and double fertilization (18–20, 22, 23, 32–37). Therefore, we used high resolution 3D live imaging of PT reception by two-photon excitation microscopy (2PEM) to analyze the penetration of the receptive synergid by the PT and associated intracellular signalling, represented by cytosolic calcium ([Ca^2+^]_cyt_) imaging. We show that, during penetration, the receptive synergid forms a membrane invagination that surrounds the invading PT, a structure we term the peri-tubular membrane (PRM).

Although the invasion process appears similar to hyphal invasion and the PRM is reminiscent of the peri-arbuscular membrane, no structure similar to the pre-penetration apparatus was observed upon arrival of the PT when using synergid-expressed cytosolic and actin markers as indicators (26). In the *fer* mutant synergids, a PRM was formed, suggesting that *FER* does not regulate entry of the PT into the receptive synergid. Finally, we demonstrate that PRM formation is concomitant with distinct cytosolic calcium ([Ca^2+^]_cyt_) signatures in synergids and PTs, and that PTs deficient in *AUTOINHIBITED Ca^2+^ ATPASE9* (*ACA9*) activity show aberrant [Ca^2+^]_cyt_ spiking and often fail to enter the receptive synergid.

## Results

### The pollen tube is enveloped by the peri-tubular membrane formed by the receptive synergid during phase II of PT reception

To discern the spatial relationship of the PT and synergids during PT reception, we first performed widefield live cell imaging using the semi-*in vitro* (SIV) method with excised ovules (35). Using the [Ca^2+^]_cyt_ sensor R-GECO1 expressed in the PT under the *pLAT52* promoter and the roGFP2-Orp1 hydrogen peroxide sensor expressed from the synergid-specific *pMYB98* promoter, we tracked the tip of the PTs and observed that in seven cases of successful PT reception, the PT grew towards the chalazal pole of the synergids before rupture (Movie S1A-I). In two cases of successful PT reception, the PT, after reaching the chalazal pole of the synergids, grew back around towards the micropylar pole before it ruptured (Movie S1 H,I), and in three instances of failed PT reception, the synergid signal rapidly spread into the neighboring central cell, suggesting rupture of both cell membranes (Movie S1J-L). Growth of the PT during successful PT reception showed three growth phases as previously reported (18), growing at a mean of 1.4 um/min during PT guidance, 0.47 um/min after arrival at the FA (phase I), and 1.66 um/min after the PT resumed rapid growth (phase II) and the PT and synergid [Ca^2+^]_cyt_ signals overlapped.

Our observations by widefield epifluorescence microscopy support previous imaging results that show an overlap of the synergid and PT signals during phase II; however, they lack the resolution necessary to distinguish whether the PT grows inside or outside of the synergids (18, 19). To address this problem, we took transgenic marker lines carrying a *pLAT52:dsRED* or *pMYB98:HyPer7-NES* construct for live imaging of PT arrival by widefield microscopy, followed by 3D reconstruction using 2PEM during the three growth phases of PT reception (Fig. 1A). In phase II, before rupture, the PT displaced the cytosol of the receptive synergid. After rupture, the PT’s cytosolic marker dsRED occupied the space of the receptive synergid and extended around the adjacent egg cell (Fig. 1B-C). In rare instances of receptive synergid rupture, the cytosol of the receptive synergid leaked into the central cell and the PT continued to grow towards the egg cell before arresting its growth (Fig. 1D-E). Together, these results suggest that the PT pushes into the receptive synergid before rupture and that, under *in vitro* conditions, aberrant penetration of the central cell sometimes occurs as the PT pushes into and through the receptive synergid.

**Figure 1.**
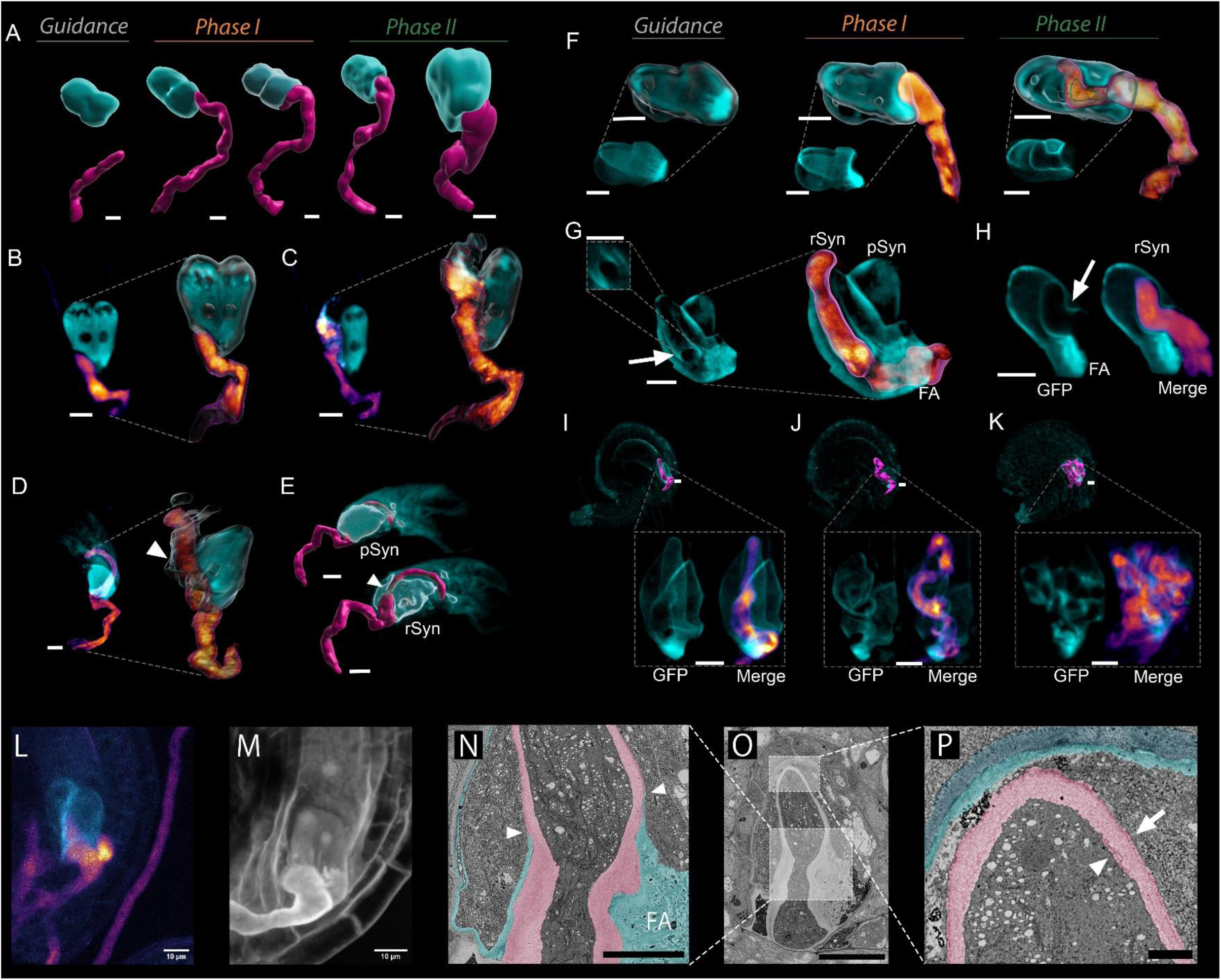
3D reconstruction of PT-synergid invasion using cytosolic and membrane-associated markers. (A) 3D reconstruction and segmentation of the cytosolic marker proteins HyPer7-NES (synergids, cyan) and dsRED (PT, magenta) in different growth phases and orientations. (B, C) Averaged signal of a Z projection (left) and enlarged 3D surface reconstruction overlayed with GFP signal (right) of the same cytosolic markers 5 minutes before (B) and after PT rupture (C). (D, E) Representative instance of receptive synergid (rSyn) rupture without PT rupture where signal could be seen inside the central cell. (D) Averaged signal of a Z projection (left) and enlarged 3D surface reconstruction of the same cytosolic markers (right) (E) Opposite sides of 3D reconstructed and segmented synergid rupture overlayed with GFP signal. Arrow heads shows site of synergid rupture. pSyn, persistent synergid. (F) Averaged signal projection of synergid membrane marker ROP2-GFP (bottom) and enlarged 3D surface reconstruction with PT cytosolic marker dsRED and overlayed with GFP and dsRED signal (top). (G) Averaged signal projection of synergid membrane marker ROP2-GFP (left) and enlarged average projection merged with PT cytosolic marker dsRED showing the penetration peg-like structure (squared), PRM (arrow), filiform apparatus (FA), receptive synergid (rSyn), and persistent synergid (pSyn). (H) Averaged signal projection of ROP2-GFP-expressing synergid section (left) and merged with the dsRED-expressing PT (right), showing the PRM (arrow) and filiform apparatus (FA). (I, J, K) Averaged signal projection of *fer-1/fer-1* mutant ovules expressing MARIS-GFP with 3D reconstructed and segmented dsRED-expressing-PT with enlarged average projection of just synergid GFP signal and merged with the PT at different stages of coiling. (L – P) Scanning electron microscopy (SEM) array tomography analysis of PT-synergid invasion. (L) Ovule expressing the fluorescent proteins FER-GFP, ROP2-GFP and pLifeAct-HyPer7 being invaded by a dsRED-expressing PT imaged live with 2PEM. (M) The same ovule as in (L) that has been fixed and imaged with SR2200 staining and 2PEM. (N-P) SEM images of ovule in (L-M) that has been sectioned. (N) Site of PT penetration showing the cell walls of the PT (pink) and synergid (cyan) as distinguished by the middle lamella and false colored for visualization. Arrowheads show where the synergid cell wall has thinned out surrounding the penetration site of the PT (scale bar is 5 μm). (O) Zoomed out SEM image of showing the relative location of the zoomed in images shown in (N, P). (P) Tip of the PT at the chalazal end of the synergid showing the cell walls of the integuments (dark blue), synergid (cyan), and PT (pink). Arrow indicates the PRM and arrowhead indicates the PT plasma membrane (Scale bar is 1μm). All images are from ovules and PTs grown semi-*in vitro* and taken by 2PEM. All scale bars are 10 μm unless indicated otherwise. Levels of images were adjusted and a gaussian filter applied to the GFP channels and smooth filter to the RFP channels.

Using 3D reconstruction with cytosolic markers could, however, not distinguish whether the PT is in or outside the receptive synergid during phase II. Thus, we performed the live imaging using two membrane-associated markers expressed in the synergids under the control of the *pMYB98* promoter: MARIS-GFP, a myristoylated cytoslolic receptor-like kinase, and ROP2-GFP, an S-acylated type II Rho GTPase. Both are cytosolic proteins that have an affinity for the plasma membrane due to their lipidation, and thus offer the advantage of simultaneously marking the cytosol and plasma membrane (38–40). Using these markers, PTs in phase I were observed to distort the FA of the receptive synergid and, in phase II, PTs were enveloped by a membrane extending from the FA (Fig. 1F-H). Because this membrane, which was not reported before, surrounds the invading PT similar to the peri-arbuscular membrane surrounding an invading mycorrhizal hyphae, we termed this structure the “peri-tubular membrane” (PRM). The PRM was also seen in *fer-1* mutants where PTs often fail to rupture, resulting in extensive distortions of the synergid membrane (Fig. 1I-K). 3D reconstruction and surface rendering of the PRM in *fer-1* mutant ovules confirmed that the PT becomes completely enveloped by the synergid membrane in *fer-1* synergids (Movie S2). Together, these results demonstrate that, both in wild-type and *fer-1* synergids, the PT penetrates the receptive synergid, but is topologically not inside the receptive synergid as it is still surrounded by the PRM, a plasma membrane of the receptive synergid, similar to mycorrhizal hyphae that are surrounded by the plasma membrane of the plant cells they penetrate.

To determine the structure of the PRM with respect to the cell wall, we fixed ovules as the PTs penetrated them and analyzed them by SEM array tomography (Fig. 1L-P). With this approach, the PT could be observed to penetrate the receptive synergid around the FA, where the synergid cell wall surrounded the invading PT and thinned out along the PT shank (Fig. 1N). The synergid cell wall was completely devoid at the PT tip where the PT cell wall was directly surrounded by the PRM and therefore sandwiched between two plasma membranes (Fig. 1P).

### Peri-tubular membrane formation coincides with a [Ca^2+^]cyt spike in the pollen tube and [Ca^2+^]cyt flooding in the synergids

We next sought to capture the live dynamics of PRM formation during PT reception using 3D 2PEM and an improved *in vitro* system to image PT reception, the SIV *cum septum* method (41). To do so, we monitored the entire volume of MARIS-GFP-expressing synergids in wild-type and *fer-1* synergids at 15 second intervals along both the dorsal-ventral and distal-proximal axis of the ovule (Movie S3). In both orientations and genetic backgrounds, the PT was observed to slowly displace the FA before rapidly growing into the receptive synergid inside an invagination of its plasma membrane. PRM formation was concomitant with the entry of the PT into the receptive synergid, similar to peri-arbuscular membrane formation during the invasion of mycorrhizal hyphae. However, no cytoplasmic column was formed preceding penetration as it occurs with formation of the pre-penetration apparatus during mycorrhization.

To confirm the absence of a pre-penetration apparatus and simultaneously monitor [Ca^2+^]_cyt_ dynamics in the PT during invasion, we crossed a marker line expressing FER-GFP and pLifeAct-HyPer7 from the *pFER* and *pFG* promoters, respectively, in the female gametophyte with a line expressing the [Ca^2+^]_cyt_ biosensor R-GECO1 in the PT. Imaging at five second intervals using our 3D 2PEM method, no major actin cytoskeleton rearrangements were observed before or after entry of the PT into the receptive synergid, further supporting the lack of a pre-penetration apparatus, which involves actin rearrangements (Movie S4B, D) (26). As previously reported, the PTs showed [Ca^2+^]_cyt_ oscillations during phase I and produced a high amplitude [Ca^2+^]_cyt_ spike immediately before speeding up their growth and penetrating the receptive synergid (Movie S4C, F) (18). Because this prominent [Ca^2+^]_cyt_ spike immediately precedes phase II, we termed it the “transition spike”, referring to the transition between phase I and phase II of PT reception.

Next, we tested whether the transition spike is associated with PRM formation by crossing a transgenic line carrying the *pLAT52:R-GECO1* construct with a line expressing the MARIS-GFP membrane-associated marker in the synergids (Fig. 2A, Movie S5). In all cases (n=8), the transition spike preceded the rapid growth of the PT (Fig. 2B, Fig. S2) and was concomitant with PRM formation. This [Ca^2+^]_cyt_ spike occurred irrespective of successful PT rupture, as it also occurred in three of eight samples where the PTs failed to rupture (Movie S6). While the amplitude of the transition spike varied among replicates, in both successful and unsuccessful PT rupture events, we found that the amplitude of the transition spike was on average 5.2-fold change higher than the [Ca^2+^]_cyt_ spike average of the preceding seven minutes of phase I (Fig. 2C, E, Movie S6). Because it is possible that this transition spike is important for the PT to change its growth rate and commit to enter the synergid and rupture, we asked whether this transition spike is missing in PTs penetrating *fer-1* synergids. Using the same settings, we performed imaging of *fer-1* mutant synergids expressing MARIS-GFP or ROP2-GFP (n=8) (Movie S7, S8). The transition spike was still present in most PTs that penetrated *fer-1* mutant synergids; however, the amplitude was significantly lower than in the wild type, with a 2.8-fold change on average (Fig. 2D-E, Fig. S3). Notably, PTs interacting with *fer-1* synergids, in comparison to PTs interacting with wild-type synergids, showed lower acceleration during the transition as they grew faster during phase I, but slower during the transition (Fig. 2F).

**Figure 2.**
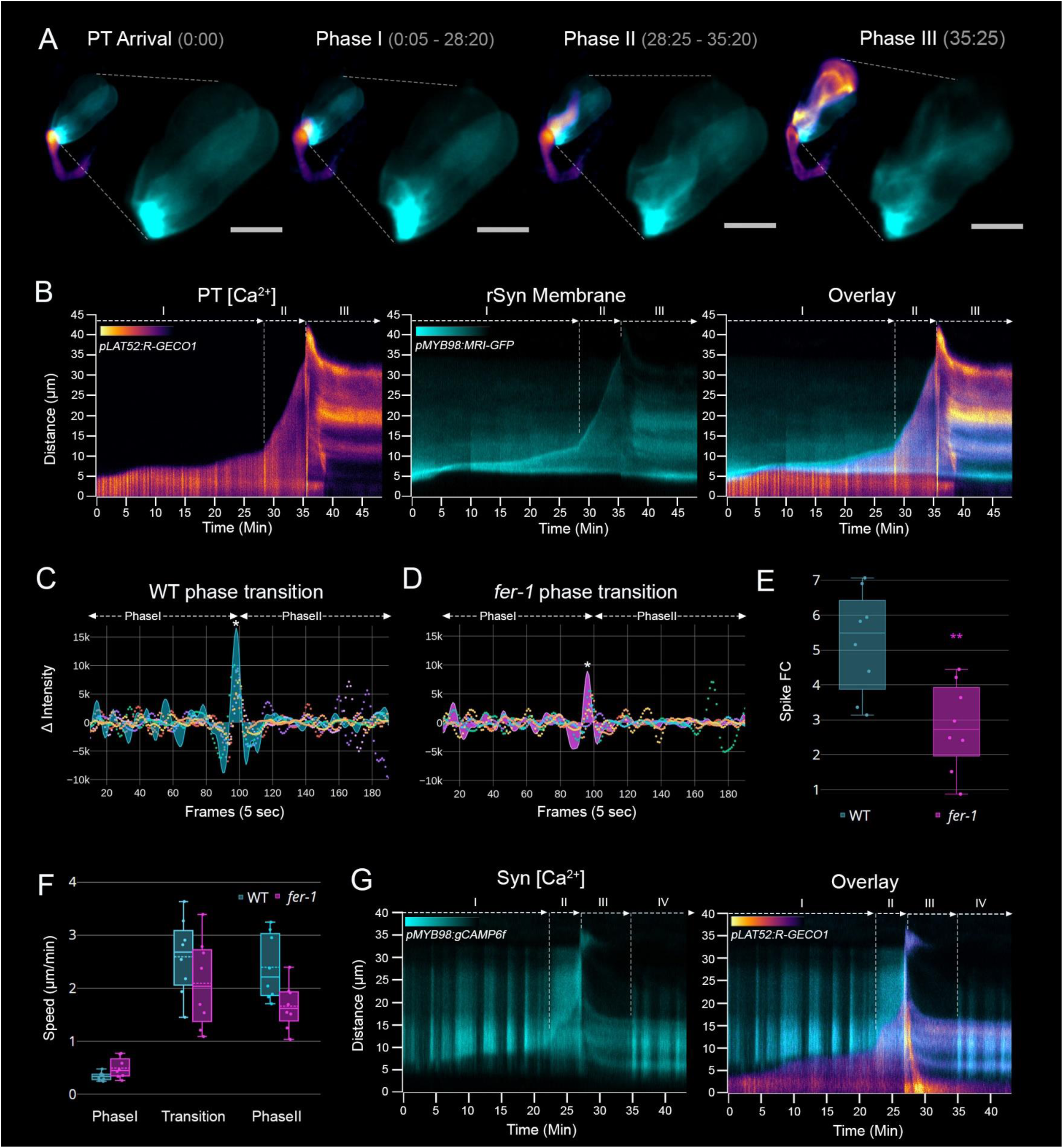
PT and synergid [Ca^2+^]_cyt_ dynamics during peri-tubular membrane formation. (A) Snapshots of live 3D imaging of a R-GECO1-expressing PT and MARIS-GFP-expressing synergids from Movie S5. Images show averaged images over the course of 4 minutes within the represented timepoints of each phase. For each phase, both the merged channels and enlarged GFP channel are shown to illustrate the synergid membrane displacement over time. Scale bar applies just for the enlarged GFP channel and is 10 μm. Time is min:sec. (B) Kymographs of live imaging shown in (A) and Movies S5. Left shows kymograph of RGECO-1 in the PT, middle shows the synergid MARIS-GFP signal, and right shows merged kymographs. Roman numerals label the different growth phases of the PT. Phase I-phase II transition is determined by the prominent [Ca^2+^] spike in the PT and rapid retreat of the synergid membrane. Phase II-phase III transition is determined by rupture of the PT and synergid degeneration. (C, D) Multicolored scatterplot of calcium oscillations from eight WT R-GECO1-expressing PTs interacting with wild-type (WT) MARIS-GFP (C) or *fer-1* MARIS-GFP and *fer-1* ROP2-GFP (D) synergids aligned around the transition [Ca^2+^] spike (asterisk) shown in Movies S5, S7. PT traces from WT ovule 6 and PT from *fer-1* ovule 3 with the largest transition [Ca^2+^] spikes are shown as a cyan line graph (C) or pink line graph (D) respectively. (E) Box plots of the transition [Ca^2+^] spike fold changes (FC) for each PT shown in (C and D) calculated as the ratio of the signal intensity of the transition [Ca^2+^] spike to the average spike maxima of the previous 7 minutes. Spike FCs between WT and *fer-1* were significantly different (p-value: 0.0037) using a t-test. (F) Box plots of the growth speed of eight WT PTs interacting with WT and *fer-1* synergids during phase I (7 minutes before phase transition), transition (1 minute surrounding transition [Ca^2+^] spike), and phase II (7 minutes following phase transition or until PT rupture). Speeds were calculated from distance travelled in the kymographs shown in Movies S5, S7. Mean WT speeds were 0.34, 2.6, and 2.4 μm/min for phase I, transition, and phase II respectively. Mean *fer-1* speeds were 0.50, 2.1, and 1.7 μm/min for phase I, transition, and phase II respectively. Solid and dashed middle lines of the box are the median and mean, respectively. (G) Kymograph of WT GCaMP6f-expressing synergids (left) and merged kymograph with R-GECO1-expressing PT (right).

In order to characterize [Ca^2+^]_cyt_ dynamics in the synergids during PRM formation at high resolution using 2PEM, we simultaneously imaged, respectively, the synergid- and PT-expressed [Ca^2+^]_cyt_ sensors GCaMP6f and R-GECO1. Consistent with previous reports, non-synchronous [Ca^2+^]_cyt_ oscillations in the two synergids were triggered by PT arrival, which were followed by [Ca^2+^]_cyt_ flooding occurring during phase II of PT growth (n=3) (Movie S9) (18, 19). As the receptive synergid’s cytosol began to be displaced by the PT in phase II, both synergids showed a sustained, global [Ca^2+^]_cyt_ influx until PT rupture (Fig. 2G). Together, these results suggest that PRM formation is concomitant with a high amplitude transition spike in the PT as it accelerates, and [Ca^2+^]_cyt_ flooding in the synergids as the plasma membrane of the receptive synergid becomes displaced.

### Mutant pollen tubes lacking AUTOINHIBITED Ca^2+^ ATPASE9 activity are deficient in phase II growth and peri-tubular membrane formation

Because PRM formation was observed even in *fer-1* synergids where intercellular signaling between the PT and synergids is defective, we asked the question whether cell invasion is a process that is actively regulated by the PT and/or synergid or simply a consequence of the PTs growth path.

To address this, we reviewed the literature for PT reception mutants showing PT growth arrest at the FA rather than *fer*-like PT coiling. The PT discharge defect of *aca9* mutant PTs reported by Schiott and coworkers (42) showed frequent growth arrest of the PTs in micropyle of pistils pollinated with *aca9* pollen. However, high resolution imaging of the PT in relationship to the synergids was not conducted nor were [Ca^2+^]_cyt_ dynamics of *aca9* mutant PTs characterized during PT growth or their interaction with the synergids. In a recent paper, the *aca9* mutant was described to have a PT overgrowth phenotype; however, the figure in the paper shows mostly ovules without *fer*-like PT overgrowth and the [Ca^2+^]_cyt_ dynamics of *aca9* PTs were not characterized (43).

Therefore, we generated CRISPR mutants in *ACA9* in marker lines carrying the *pLAT52:dsRED* and *pLAT52:R-GECO1* and isolated homozygous *aca9* mutants in the T2 generation with single nucleotide insertions in exons 18 and 30, respectively (Fig. 3A). Consistent with previous findings (42) homozygous *aca9* mutants showed a strong reduction in the percentage of fertilized ovules, with siliques of *aca9, pLAT52:R-GECO1* lines having, on average, only 29% fertilized ovules in the top and 2% in the lower half of the siliques, indicating an impairment of PT growth through the transmitting tract (Fig. 3B, C).

**Figure 3.**
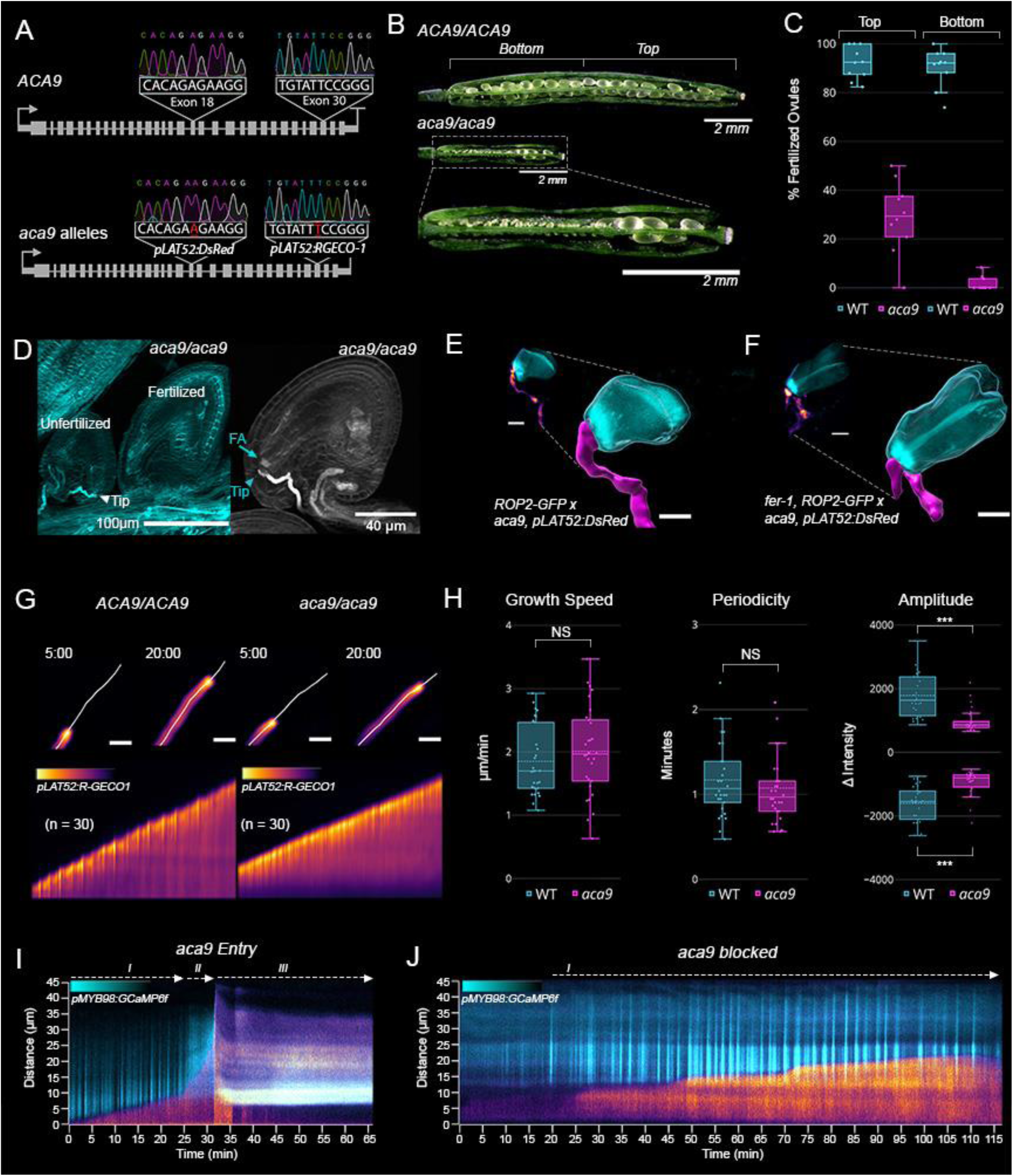
*aca9* PTs are deficient in phase II growth and peri-tubular membrane formation. (A) Gene model of *ACA9* showing the CRISPR alleles used in this study. Sanger sequencing results of 11 base pairs surrounding gRNA1 in exon 18 or gRNA3 in exon 30 in wild-type (WT) (top) and mutant lines (bottom) are shown. The mutation in the *pLAT52:dsRED* background is a single adenine insertion 3bp from the PAM site in exon 18 and the mutation in the *pLAT52:R-GECO1* background is a single thymine insertion 3bp from the PAM site in exon 30. (B) Representative siliques from WT *pLAT52:R-GECO1* and *aca9, pLA52:RGECO-1* lines dissected open to show fertilized and unfertilized ovules. Both siliques are from the fourth flower down from the first stage 16 flower. (C) Seed set of WT *pLAT52:RGECO-1* and *aca9, pLA52:RGECO-1* lines measured by percentage of fertilized ovules in either the top or bottom half of the silique as shown in panel B. (D) Representative ovules of the *aca9, pLAT52:RGECO-1* line stained with aniline blue. The cyan image shows an example of a blocked *aca9* PT at the filiform apparatus (FA) of an unfertilized ovule (left) and a ruptured *aca9* PT in the micropyle of a fertilized ovule. The gray image shows a blocked *aca9* PT, which has arrested growth at the FA. Arrowhead labelled, “Tip” and arrow labeled “FA” shows the tip of the growth arrested PT and FA, respectively. (E, F) *In vivo* imaging of dissected pistils from either WT *pMYB98:ROP2-GFP* (E) or *fer-1*, *pMYB98:ROP2-GFP* (F) 20-24 hours after pollination with *aca9, pLAT52:dsRED* PTs. Averaged images of blocked *aca9* PTs (left) and their enlarged 3D segmented reconstruction (right) are shown. Scale bars are 10 μm. (G) Representative images and kymographs of *in vitro* grown WT *pLAT52:R-GECO1* (left) and *aca9, pLAT52:R-GECO1* PTs (right). Time points at 5:00 and 20:00 (min:sec) with the line used to make the kymograph below are shown. Scale bar is 10 μm and only applies for the top PT images. (H) Box plots of growth speeds, periodicity and amplitude of 30 *in vitro* grown WT *pLAT52:R-GECO1* (cyan) and *aca9 pLAT52:R-GECO1* PTs (magenta). The mean growth speed for WT and *aca9* PTs was 1.85 and 2 μm/min, respectively. The mean period of [Ca^2+^] oscillations in WT and *aca9* PTs was 70 and 65 seconds, respectively. The mean maxima and minima amplitude for WT PTs was 1786 AU and -1602 AU respectively, which was significantly different from *aca9* mean maxima, 979 AU (p-value < 0.001) and minima, -934 AU (p-value < 0.001) . (I, J) Overlayed kymograph of *pMYB98:GCaMP6f* and *aca9, pLAT52:R-GECO1* showing WT-like entry and rupture of the PT (I) and blocked entry (J).

Aniline blue staining of self-pollinated *aca9, pLAT52:R-GECO1* siliques revealed that many PTs indeed arrive and arrest growth in the micropyle. High resolution imaging of these pistils by 2PEM showed that the tip of these PTs were arrested just at or around the FA (Fig. 3D, Fig. S4). To test whether *aca9* PTs fail to enter the synergids, we crossed *pMYB98:ROP2-GFP* pistils with *aca9, pLAT52:dsRED* pollen and dissected them 22-24 hours after pollination. All of the *aca9* PTs that had arrived at the synergids but had not ruptured, were arrested at the FA and showed no signs of PRM formation (Fig. 3E). This growth arrest was independent of *FER* as the same experiment done with *fer-1* mutant synergids expressing, ROP2-GFP frequently showed arrested PTs at the FA even in cases where several PTs entered a micropyle (Fig. 3F, Fig. S5).

To test if a difference in calcium signaling may underlie growth arrest of *aca9* PTs, we first measured the [Ca^2+^]_cyt_ dynamics of wild-type and *aca9* PTs expressing R-GECO1 in the same genetic background during *in vitro* growth. 30 wild-type and mutant PTs were imaged every five seconds on a widefield microscope and the intensity of R-GECO1 fluorescence was measured at their tip (Fig. 3G). Surprisingly, *aca9* PTs showed no significant difference in growth speed or [Ca^2+^]_cyt_ oscillation period compared to the wild type; however, the amplitude of the spikes was significantly reduced (Fig. 3H). We next analyzed the [Ca^2+^]_cyt_ dynamics in *aca9* PTs during PT reception by the synergids using 2PEM with the same settings as for imaging R-GECO1 in the wild type. Of the eight PTs imaged, five entered the synergid with four of them successfully rupturing, while three were arrested at the FA (Fig. S6, Movie S10, S11). As we had observed during *in vitro* growth, the spike amplitude while signaling with the synergids was also significantly reduced compared to the wild type (Fig. S7). When imaging the [Ca^2+^]_cyt_ sensor *GCaMP6f* in the synergids using the semi-*in vitro cum septum* method, *aca9* PTs that behaved like the wild type could also initiate phase I [Ca^2+^]_cyt_ oscillations and phase II [Ca^2+^]_cyt_ flooding in the synergids. In the case of blocked *aca9* PTs, phase I [Ca^2+^]_cyt_ oscillations were initiated in the synergids upon PT arrival and persisted for over 2 hours without PT entry (Movie S12). Together, these results suggest that *aca9* PTs have a synergid entry phenotype with partial penetrance, and lower amplitude of [Ca^2+^]_cyt_ spiking during PT growth and signaling with the synergids. Importantly, the previously reported PT discharge defects of *aca9* PTs can be attributed to a failure to enter the synergids, as those PTs that entered and formed a PRM and ruptured similar to the wild type.

## Discussion

Our study on synergid invasion by the PT using a live 3D 2PEM imaging approach has clarified the spatial relationship of the PT and synergids during PT reception and confirmed that the PT penetrates the receptive synergid during phase II. Using this approach, we found a previously undescribed structure, the PRM, a membrane invagination of the synergid’s plasma membrane that rapidly forms upon PT entry and envelopes the PT during its rapid growth prior to rupture. By identifying this new structure and providing the highest resolution videos of PT reception to date, this study provides a better understanding of this complex process and facilitates the formulation of more accurate models of the signaling mechanism leading to sperm release. For example, we observe that PT rupture only occurs after penetrating the synergid cell wall and forming the PRM. Therefore, the signal(s) triggering PT rupture must only be produced or perceived after the entrance of the PT, providing a means by which the receptive synergid can control the timing and orientation of sperm release. This understanding must be integrated into current models of PT rupture, which propose that it is mediated by the perception of RALFs secreted by the ovule (44) that, however, would purport PT rupture outside of the synergids. Further research on how the engulfment of the PT by the PRM affects the integrity of the PT’s cell wall and leads to cell death should shed light on how the synergids control PT rupture.

In addition to the synergids expressing several genes that are also important for interactions with fungi, the newly discovered PRM being reminiscent of the peri-arbuscular membrane provides yet further parallels between PT reception and invasion by fungal hyphae. Still, there are distinct differences. For example, invasion of epidermal and cortical root cells by mycorrhizal fungi occurs over a much longer time scale and is accompanied by large cellular rearrangements including nuclear migration, cytoskeletal restructuring, lipid synthesis, and construction of a cytoplasmic column during the formation of the pre-penetration apparatus (26). In contrast, PT reception occurs over the course of only about one hour and we found no evidence to support the formation of a pre-penetration apparatus upon PT arrival. Instead, the extensive plasma membrane invaginations of the FA may pre-emptively prepare the synergids for invasion by the PT since they are destined to be penetrated, unlike root epidermal cells of which only a few will interact with mycorrhizal hyphae. Thus, we propose that, in addition to previously its described functions, the FA itself acts as a pre-penetration apparatus and, because it is already formed prior to PT arrival, allows for the rapid entry of the PT into the synergid. Another distinction relates to calcium signaling during interaction with mycorrhizal fungi, which is mostly occurring in the nuclear periplasm where calcium plays a role in the transcriptional reprogramming of the cell. In contrast, calcium signaling in the synergids has only been shown in the cytoplasm and due to the short duration of PT reception, it is unlikely that there is major transcriptional reprogramming of the synergid upon PT arrival.

Our calcium imaging showed that the transition of PT growth from phase I to phase II is concomitant with a high amplitude [Ca^2+^]_cyt_ spike in the PT and [Ca^2+^]_cyt_ flooding in the synergids. These calcium signatures are associated with PRM formation and possibly the result of abrupt changes in forces acting on the PT and the synergid plasma membranes as the PT rapidly accelerates when growing into the synergid. By generating *aca9* mutations in PT marker lines, we showed that this calcium pump is important for entering the receptive synergid *in vivo*. Rejection of *aca9* PTs is unlikely to be regulated by the synergids as *aca9* PTs still trigger [Ca^2+^]_cyt_ oscillations in the synergid and arrest growth at the FA of *fer* mutant synergids, where synergid [Ca^2+^]_cyt_ oscillations are impaired. However, the lower amplitude [Ca^2+^]_cyt_ spiking in *aca9* PTs suggests that calcium signalling during phase I may actively regulate synergid entry. But it cannot be excluded that some *aca9* PTs may simply accumulate cytotoxic levels of calcium, causing them to arrest growth at the FA after several hours of growth. Future studies focused on local changes in the synergid and PT cell walls as the PT breaks into the synergid should help to better understand the phenotype of *aca9* PTs and address whether localized synergid cell wall degradation is mediated by the PT or the synergids.

## Materials and Methods

### Transgenic Plant Generation

#### Gateway cloning

The construct *pMYB98:MARIS-GFP* was generated by gateway cloning. The MYB98 promoter was PCR amplified from pLM4 (45) using primers LM218 (5′-CCCTCTAGATGTTTTGGAAAGGAGAAAAAA-3′) and LM219 (5′-TTTAAGCTTATACACTCATTGTCCTTCG-3′) introducing the HINDIII and XbaI restriction site. This product was then column purified and ligated to pMDC83 (46) that was cut with HindIII and XbaI, desphosphorylated with phosphatase enzyme, and purified from a gel to yield pNJD15. This destination vector was then used for an LR reaction with entry vectors ABD43 (MARIS CDS without a stop codon in pDONR207) (38). For *pMYB98:ROP2-GFP,* the ROP2 CDS without stop codon was PCR amplified from flower cDNA with primers NJD50 (5′-GGGACAAGTTTGTACAAAAAAGCAGGCTTAATGGCGTCAAGGTTTATAAAGTG-3′) and NJD51 (5′-GGGGACCACTTTGTACAAGAAAGCTGGGTACAAGAACGCGCAACGGTTC-3′), purified from a gel, and inserted into pDONR221 with a BP reaction to yield the entry vector pNJD28. A LR reaction between pNJD28 and pNJD15 yielded *pMYB98:ROP2-GFP* (pNJD33). *pMYB98:roGFP2-Orp1* was made by an LR reaction between *roGFP2-Orp1* in pDONR207 (47) and the LM4 destination vector (45).

#### GreenGate Cloning

*pMYB98:GCaMP6f* was generated using GreenGate cloning with modified protocols (48). For the *MYB98* promoter module (pGGA-Myb98), 713 bp upstream of the START-codon of *MYB98* (At4G18770) was PCR-amplified from gDNA (Fwd: 5′-ACAGGTCTCAACCTACACTCATTGTCCTTCGGCA-3′; Rev: 5′-ACAGGTCTCTTGTTCATTGTTTTGGAAAGGAGAA-3′) and cloned into pGGA000. To generate the expression construct, entry vector modules pGGA-Myb98, pGGB003 (B-Decoy), pGGC-GCaMP6f (49), pGGD002 (D-Decoy), pGGE009 (Ubi10-Terminator), and pGGF009 (BastaR) were fused into the destination vector pGGZ003.

#### CRISPR cloning

The *ACA9* CRISPR constructs were cloned by phosphorylating and annealing the guide RNA primer pair (Fwd: 5′-ATTGAGACAGAATAGCCTTCTCTG-3′; Rev: 5′-AAACCAGAGAAGGCTATTCTGTCT-3′) targeting *ACA9* exon 18 and guide RNA primer pair (Fwd: 5′-ATTGGATGAAATGAATGTATTCCG-3′; Rev: 5′-AAACCGGAATACATTCATTTCATC-3′) targeting *ACA9* exon 30 with T4 Polynucleotide Kinase and cool down thermocycle from 95°C to 25°C at -5°C/min. The annealed primers contain overhangs that were ligated to the CRISPR pKAMA-ITACHI vector, pKIR1.1, that was digested with restriction enzyme AarI (50).

#### Golden Gate Cloning

The coding DNA sequences (CDSs) of HyPer7-NES and pLifeAct-HyPer7 were amplified and modified from plasmids obtained through www.addgene.org (plasmid numbers: 22692, 136467 and 136464 respectively). To perform Golden Gate cloning, we silenced the BsaI sites in the CDS of HyPer7-NES and added them in the linker regions through PCR amplification with the following primers: MB7 (Fwd: 5′ GGTCTCTAATGCACCTGGCTAATGAGGAG-3′), MB18 (Rev: 5′ AAGGGTGTTGGTGAGTCCCTG -3′), MB17 (FWD: 5′-ACAGCGCATTCAGGGACTCAC-3′), MB8 (Rev: 5′-GGTCTCTCGAATGACAACAACAATTGCATTC-3′).

We silenced the BsaI sites in the CDS of pLifeAct-HyPer7 with the following primers: MB5 (Fwd: 5′-GGTCTCTAATGGGCGTGGCCGACTTGATC-3′), MB16 (Rev: 5′-AAGGGTGTTGGTGAGTCCCTG -3′),MB15 (Fwd: 5′-ACAGCGCATTCAGGGACTCAC-3′), and MB6 (Rev: 5′-GGTCTCTCGAATGACAACAACAATTGCATTC-3′).

Level 0 CDS regions were inserted in miniT2.0 plasmids with the NEB PCR cloning kit. For the *MYB98* promoter, 1669bp upstream the gene *MYB98* has been considered and amplified using the golden gate compatible primers STB122 (5’-GGTCTCAGGAGATAGTGGCTGAGAGGT -3’) and STB123 (5’-GGTCTCCCATTGTTTTGGAAAGGAGA-3’). For the *FG* promoter, 2953 bp upstream of the gene At5G62150 has been amplified in two parts to domesticate the BsaI site inside the region of interest. The first part using the primers STB6 (5’-GTGGTCTCAGGAGTTTCTCTCACTATTAACAATTG-3’) and STB7 (5’-GTGGTCTCTCACTTGTGCCTTGCGGTGA-3’), and the second part with the primers STB8 (5’-GTGGTCTCAAGTGACCAAGTCTCTAACC-3’) and STB9 (5’-GTGGTCTCTCATTGCCTTTAATTTATAAAA-3’). The CDS and promoters were then assembled using the golden gate system.

#### Transformation and Selection

All constructs were electroporated into GV3101 agrobacterium and transformed into Arabidopsis using the floral dip method (51). All of the gateway and golden gate cloned constructed were transformed into Ler0, *fer-1/+,* or *fer-1/fer-1* plants and the transgene and *fer-1* mutation homozygosed in the T2 generation. The plasmid *pFER:MARIS[R240C]-GFP* was gifted by Aurélien Boisson-Dernier and contains a dominant active version of MARIS that was transformed into *fer-1/fer-1*. This line was used for the images in Figure 1 (I-K) and Figure S2 where it is simply referred to as *pFER:MARIS-GFP* to not confuse the reader. The GreenGate *pMYB98:GCaMP6f* construct was transformed into Col-0 and homozygosed in the T2 generation. The CRISPR vectors were transformed into both *pLAT52:R-GECO1* and *pLAT52:dsRED* backgrounds and selected for red seed fluorescence. Homozygous mutants were identified by sequencing in the T1 and the CRISPR cassette was segregated out by selecting non fluorescent seeds in the T2 where the mutations were again confirmed by sequencing. The *pLAT52:R-GECO1* and *pLAT52:dsRED* lines used were previously described (18, 21).

### Microscopy

#### Widefield imaging

For the widefield PT tracking experiment, the semi*-in vitro* (SIV) PT reception method with excised ovules was performed as first described in (35). Time-lapses were started 1 hour after donor stigmas were pollinated with pollen expressing R-GECO1 and dissected onto pollen germination medium (52). The dish was placed in the humidity chamber on the microscope and imaged using a 10X objective lens every 60 seconds for up to 9 hours with four fluorescent channels: Channel (1) 560/640 (ex/em) for R-GECO1 (25 ms), Channel (2) 488/520 for roGFP2-Orp1 (200 ms), Channel (3) 387/520 for roGFP2-Orp1 (200 ms), and Channel (4) 387/640 for background subtraction (200 ms). An autofocus function using a 100 mM range with 7 mm coarse stepping and 1 mm fine stepping was employed each minute before image acquisition. For calcium imaging of WT and *aca9* PT growth *in vitro*, the same setup was used without any ovules or autofocus and imaging using just Channel (1) 560/640 for R-GECO (25 ms) every 5 seconds for 20 minutes.

#### Live 2PEM imaging

All calcium live imaging experiments with 2 photon excitation microscopy (2PEM) were performed using the SIV *cum septum* method (41) and an inverted Leica SP8 multiphoton microscope. Calcium imaging with 2PEM was performed using 1020nm excitation at 2% power and detection with the hybrid detectors HyD1 (525/50nm) for GFP/ GCaMP6f and HyD4 (585/40nm) for R-GECO1. A Fluotar VISIR 25x/0.95NA water objective was used at 7.5 digital zoom, 1,8000 Hz scan speed for 62µm physical length in X and Y and 12 slices in Z for a total of 22µm with sampling every 5 seconds. A 19 step autofocus function was performed every 40th stack using HyD1 with a 80µm range to correct for sample drift in Z. For 3D surface reconstructed images, stacks were also taken with using a Fluotar VISIR 25x/0.95NA water objective on the Leica SP8 multiphoton microscope with 200nm – 600nm voxel size in Z. a*ca9* pollination experiments were performed by dissecting pMYB98:MARIS-GFP or pMYB98:ROP2-GFP, *fer-1* pistils 22-24 hours after pollination with *aca9* or WT *pLAT52:dsRED* pollen. Synergids were assessed for PRM formation and imaged using excitation at 960nm and 1040nm and detection with HyD1 and HyD4.

#### Aniline blue imaging

Aniline blue staining was performed as described previously and imaged with 2PEM was excited at 800nm and detected using HyD1 (525/50nm) with 0.5um Z stack steps (21).

#### Correlative light and electron microscopy

Ovules expressing FER-GFP, ROP2-GFP, and pLifeAct-HyPer7 were observed with 2PEM for penetration by dsRED-expressing PTs grown using the SIV *excised ovules* method (35). At the time of synergid penetration, a Z-stack was taken and the samples were fixed with a fixative solution containing 4% formaldehyde and 2% glutaraldehyde in cacodylate buffer (0.1 M pH: 7.4) and lightly infiltrated under vacuum. Samples were stored overnight at 4 °C before replacing the fixative solution with 0.1% (v/v) SCRI Renaissance 2200 (SR220) staining solution in cacodylate buffer for 2PEM imaging of the fixed tissue using 850nm excitation at 4.7% power and detection with hybrid detector HyD2 (460/50nm) for 250, 200nm Z slices. The staining solution was then replaced with the fixative solution and samples processed for electron microscopy.

Briefly, samples were fixed with 1% OsO_4_ for 1 hour in 0.1 M cacodylate buffer at 0°C, and 1% aqueous uranyl acetate for 1 hour at 4°C. Samples then were dehydrated in an ethanol series and embedded in Epon/Araldite (Sigma-Aldrich): 66% in propylene oxide overnight, 100% for 1 h at RT and polymerized at 60 °C for 20 h . Ultrathin serial sections (70nm) were collected on silicon wafers using an ultramicrotome (Artos 3D, Leica Microsystems) equipped with a silicon wafer holder (CD-FH, Germany, (53)). Sections were imaged in an Apreo 2 VS scanning electron microscope using the MAPS software package for automatic serial section recognition and image acquisition (Thermo Fisher Scientific, (54)). The following parameters were used for imaging: OptiPlan mode, T1 detector; pixel size of 7 nm, pixel dwell time of 3 or 5 µs, an electron high tension of 1.8 keV, and a beam current of 0.1 nA. Serial section tiff images were aligned using the FIJI71-plugin TrakEM2 (55, 56).

### Image Processing

#### PT tracking

For the widefield tracking experiment, circular rois were manually placed at the PT tip and the synergid cells were tracked by thresholding the signal and using analyze particles in Fiji. ROIs and cell outlines were flattened into the final videos and combined together in Fiji. For manual tracking of the PT tip with 2PEM calcium imaging, circular rois were placed manually at the tip and interpolated in Fiji. For videos with substantial drift, the plugin, “MultiStackReg” was employed for registration.

#### 3D reconstruction

3D reconstructions were performed using surface rendering in IMARIS after post-acquisition computational clearing using Lecia’s Thunder package. Reconstructions were segmented by signal intensity and trimmed to show the synergids (cyan LUT) and the PT (fire or magenta LUT). Transparent surfaces were overlayed with the channels to highlight the surfaces and signal intensity in panels such as Figure 1B. Panels were arranged and annotated in Adobe Photoshop CC version 24.6 (Adobe, https://www.adobe.com/).

#### Calcium measurements

All kymographs were made using a segmented line and the Multi Kymograph function in Fiji with line width set to 7. Calcium oscillation measurements were performed by converting kymographs of raw images to text images and analyzed with the data analysis pipeline ’Computational Heuristics for Understanding Kymographs and aNalysis of Oscillations Relying on Regression and Improved Statistics’, or CHUKNORRIS in “R studio” (57). For the *in-vitro* growth analysis of *aca9, pLAT52:RGECO* tip growth speed and period of oscillation was taken from CHUKNORRIS and tip fluorescence was analyzed for amplitude in MATLAB using the “find peaks” function with minimum peak distance set to 3, minimum peak prominence set to 1000, and minimum peak height set to 300. For PT calcium measurements in the PT reception assay with 2PEM, raw stacks were SUM projected in Fiji and the PT tip manually tracked as described above. The amplitude for the calcium measurements in the PT reception assay with 2PEM was performed by using the time series analysis module in CHUKNORRIS of the PT tip fluorescence taken from phase I growth and “find peaks” function in MATLAB with minimum peak distance set to 3, minimum peak prominence set to 1000, and minimum peak height set to 100. For phase transition measurements, the videos were cropped to 200 frames aligned around the phase transition at frame 100. Changes in PT speed were calculated as distance traveled in the first 95 frames (Phase I), distance traveled in frames 96-105 (Phase transition), and distance traveled in frame 106-200 or 106-rupture (phase II). For transition spike fold change calculations, the manually tracked tip fluorescence for the 200 frames was analyzed by the time series analysis module in CHUKNORRIS and amplitudes determined in MATLAB as described above. Transition spike fold change was calculated as the transition spike intensity divided by the average of the phase I spikes in the preceding 95 frames. All plots were made in Plotly Chart Studio (https://chart-studio.plotly.com/).

## Acknowledgments

We thank Celia Baroux (University of Zurich) for advice and helpful feedback about microscopy and experimental design; Aurélien Boisson-Dernier (Université Côte d’Azur, INRAE, CNRS, Institut Sophia Agrobiotech) for donating plasmids; Hannes Vogler (University of Zurich) for advice on data analysis, Valeria Gagliardini, Arturo Bolaños, Peter Kopf, Daniela Guthörl, and Frédérique Pasquer (University of Zurich) for general lab support; and Christof Eichenberger (University of Zurich) for advice and help with microscopy. We are indebted to Jose Maria Mateos Melero, Andres Käch, and Urs Ziegler (Center for Microscopy and Image Analysis, University of Zurich) for help with the SEM studies.

## Author Contributions

U.G. raised funds and supervised the project; N.D. and U.G. conceived the study; N.D. designed and performed all experiments; N.D processed and analyzed all the data; N.D. cloned and generated the transgenic lines; P.D. provided the *pMYB98:GCaMP6f* line, M.B. cloned the HyPer7-NES and pLifeAct-HyPer7 entry vectors, S.B. cloned the *pMYB98* and *pFG* promoters for Golden Gate cloning. N.D and U.G. wrote the manuscript; all authors commented on the manuscript.

