## Supplemental Figures and Movie Legends for "The Pollen Tube Penetrates the Synergid Cell by Formation of a Peritubular Membrane"

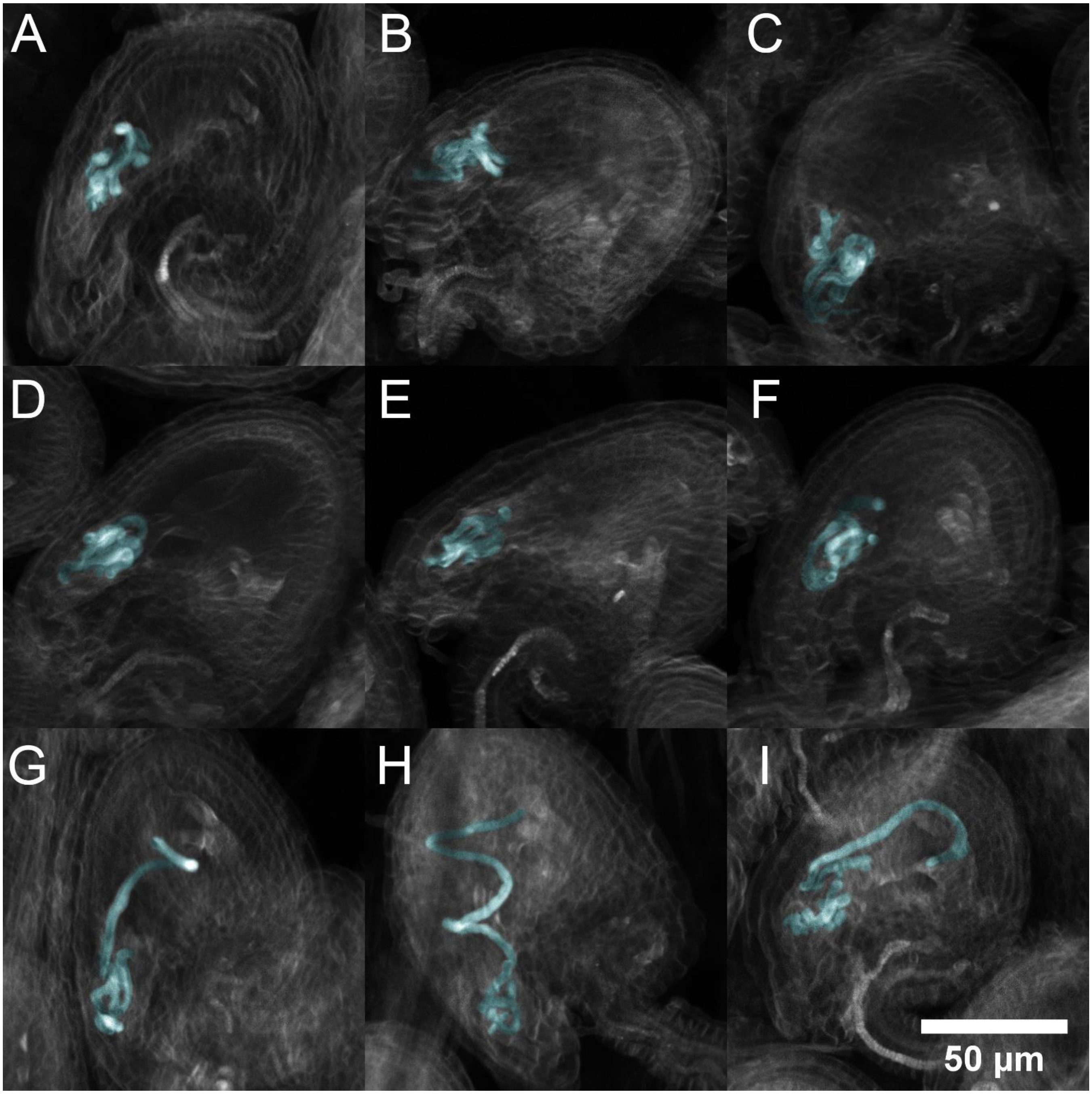


**Supplemental Figure 1. Pollen tube overgrowth phenotype in *fer-1* ovules.**

(I - J) Average projections of Aniline blue stained PTs overgrowing in *fer-1* ovules imaged using 2PEM. PTs are false colored in cyan. Scale bar is 50 µm.

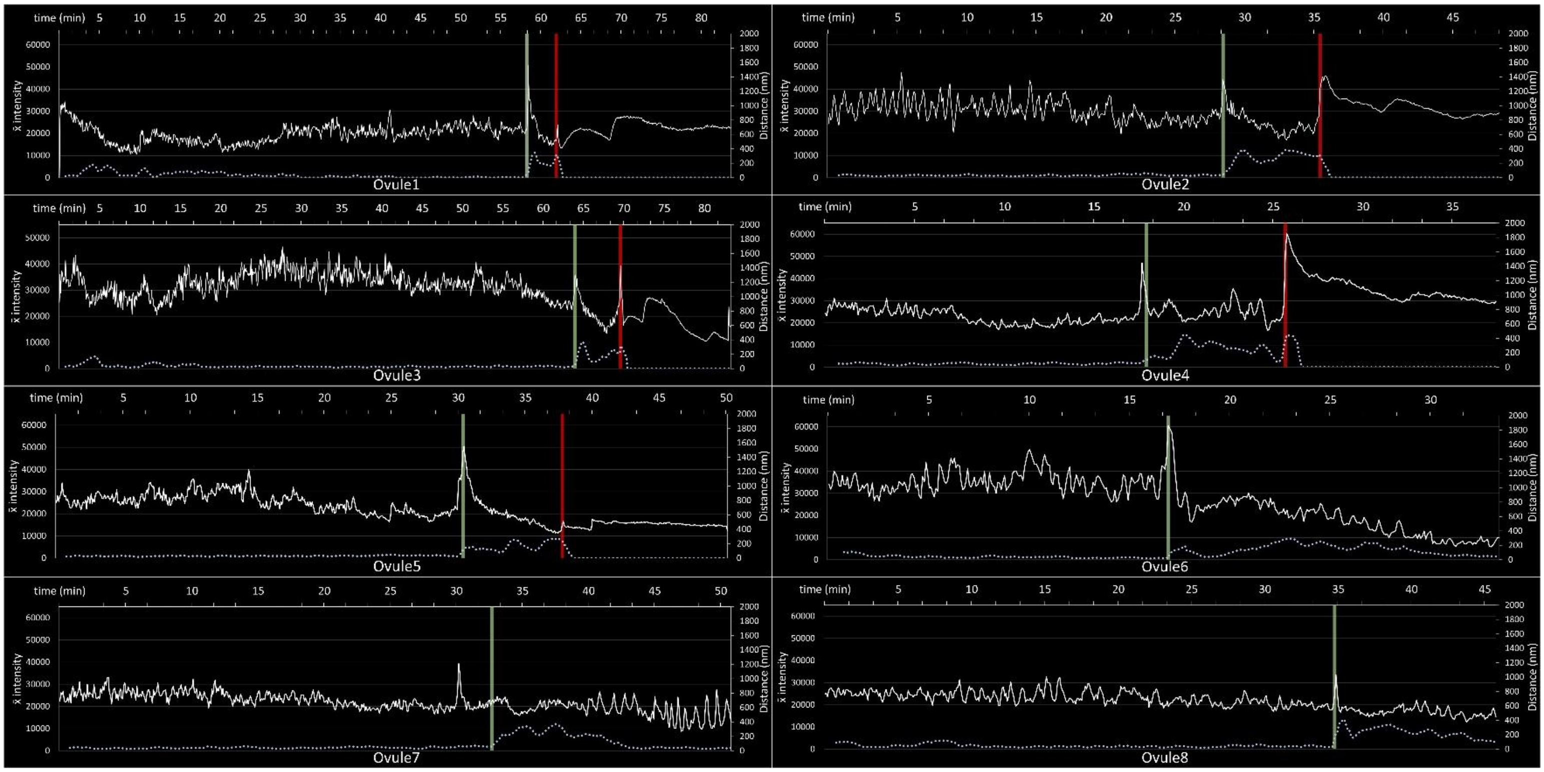


**Supplemental Figure 2. WT PT R-GECO1 $[Ca^{2+}]_{cyt}$ traces during PT reception.**

Eight $[Ca^{2+}]$ traces of manual tip tracked R-GECO1-expressing PTs interacting with *pMYB98:MARIS-GFP* synergids imaged every 5 seconds. The left Y axis is the mean intensity of summed projections of 16bit Z stacks and applies to the white line. The right Y axis is distance in nanometers and applies to the blue dotted line, which shows the moving average trendline (period 10) of the PT distance traveled calculated from the coordinates of the tip center of mass. The top X axis is time in minutes. The green line denotes the approximate transition point of the PT from phase I to phase II growth and the red line denotes rupture of the PT.

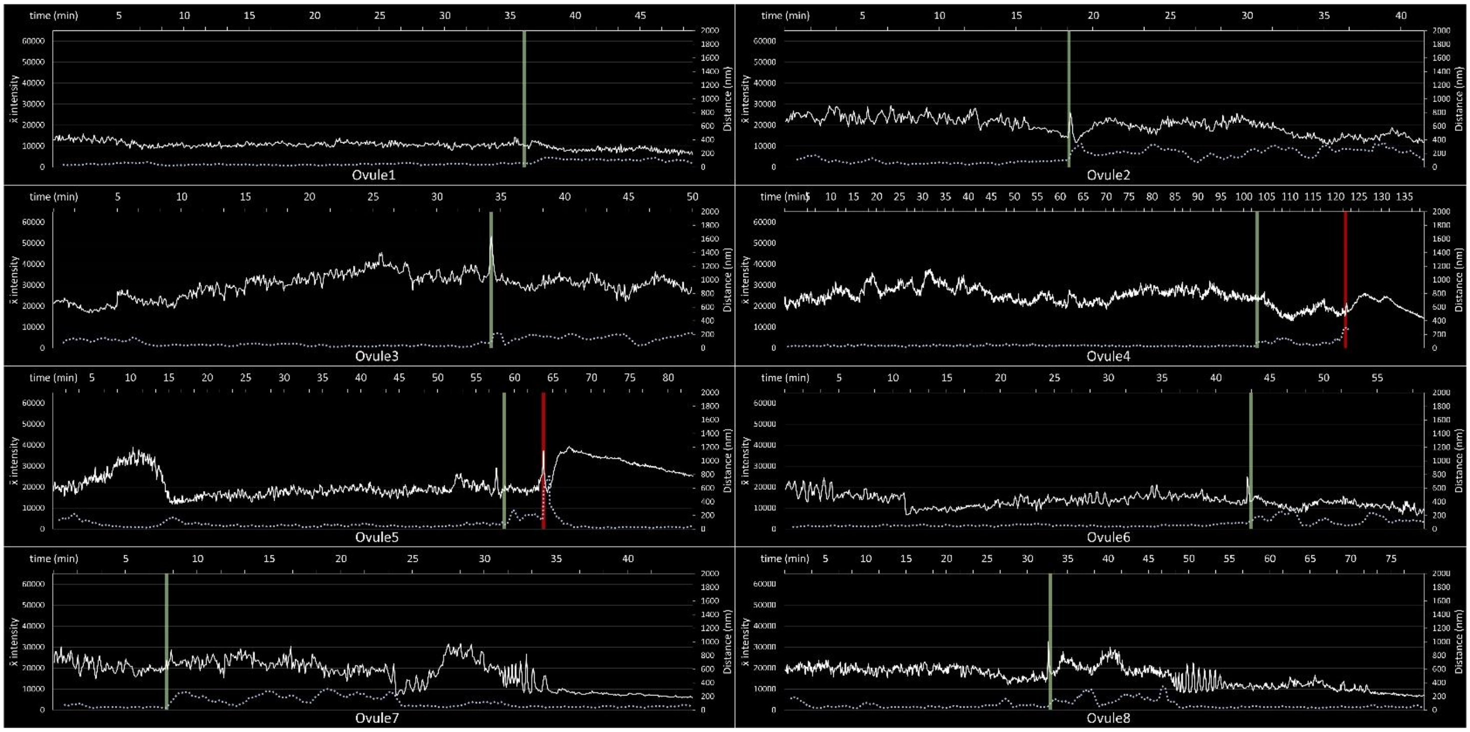


**Supplemental Figure 3. WT PT R-GECO1 $[Ca^{2+}]_{cyt}$ traces during PT reception with *fer-1* ovules .**

Eight $[Ca^{2+}]$ traces of manual tip tracked R-GECO1-expressing PTs interacting with *fer-1*, *pMYB98:MARIS-GFP* or *fer-1, pMYB98:ROP2-GFP* synergids imaged every 5 seconds. The left Y axis is the mean intensity of summed projections of 16bit Z stacks and applies to the white line. The right Y axis is distance in nanometers and applies to the blue dotted line, which shows the moving average trendline (period 10) of the PT distance traveled calculated from the coordinates of the tip center of mass. The top X axis is time in minutes. The green line denotes the approximate transition point of the PT from phase I to phase II growth and the red line denotes rupture of the PT.

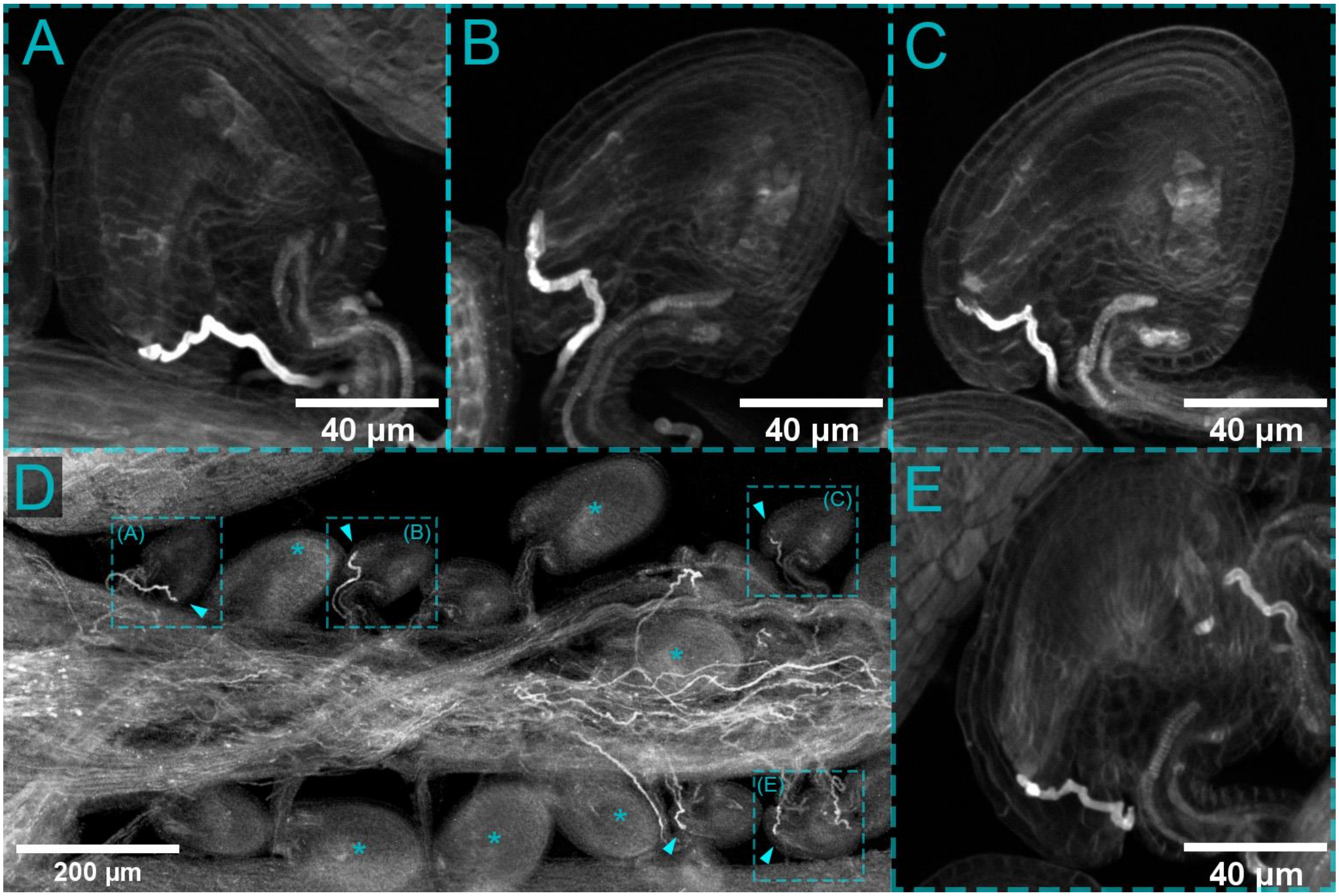


**Supplemental Figure 4. Aniline blue of *aca9/aca9* self-pollinated pistil.**

(A, B, C, E) Average projections of aniline blue stained *aca9* PTs with arrested growth at the FA in unfertilized ovules shown with dashed squares and arrowheads in (D).

(D) Merged tile scan of aniline blue stained *aca9/aca9* self-pollinated pistil. Asterisks show fertilized enlarged ovules and arrowheads show unfertilized ovules with growth arrested PTs. Boxed ovules with letters correspond to ovules shown in panels (A, B, C, E) that were imaged at higher resolution.

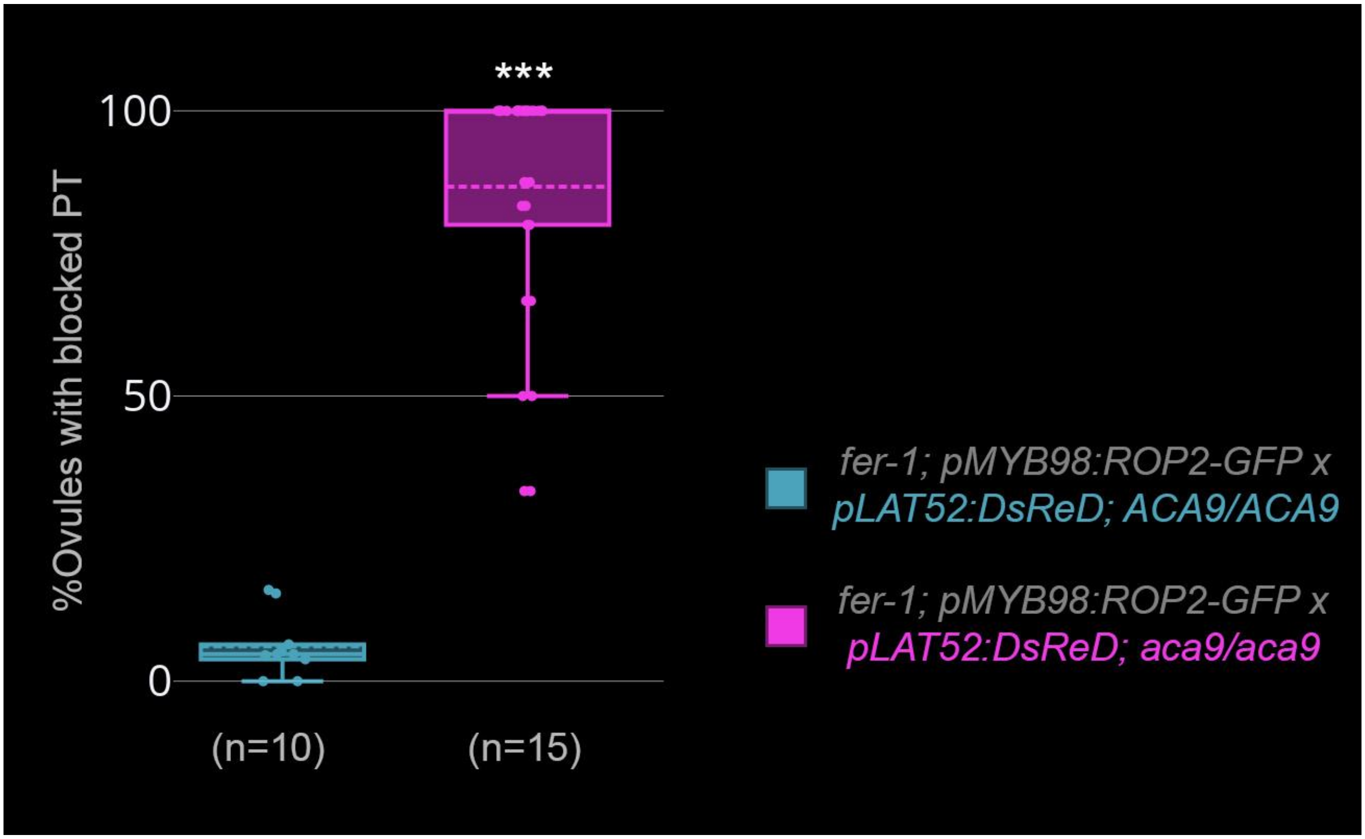


**Supplemental Figure 5.** ***aca9*** **penetration phenotype is independent of** ***FER*****.**

Box plots of the percent of *fer-1* ovules which have received a pollen tube (PT) with failure to penetrate the synergid. *fer-1/fer-1; pMYB98:ROP2-GFP* plants were pollinated with either *ACA9/ACA9; pLAT52:DsRed* (n=10 pistils) or *aca9/aca9; pLAT52:DsRed* (n=15 pistils) pollen and dissected 24 hours later to score for PT rupture, penetration without rupture (overgrowth), or failure to penetrate (blocked). Pistils with *ACA9/ACA9; pLAT52:DsRed* pollen had ovules with an average of 10% rupture, 84% overgrowth, and 6% blocked, which was significantly different (p-value < 0.001, two-sample T-test) from pistils with *aca9/aca9; pLAT52:DsRed* pollen that had ovules with an average of 0% rupture, 13% overgrowth, and 87% blocked.

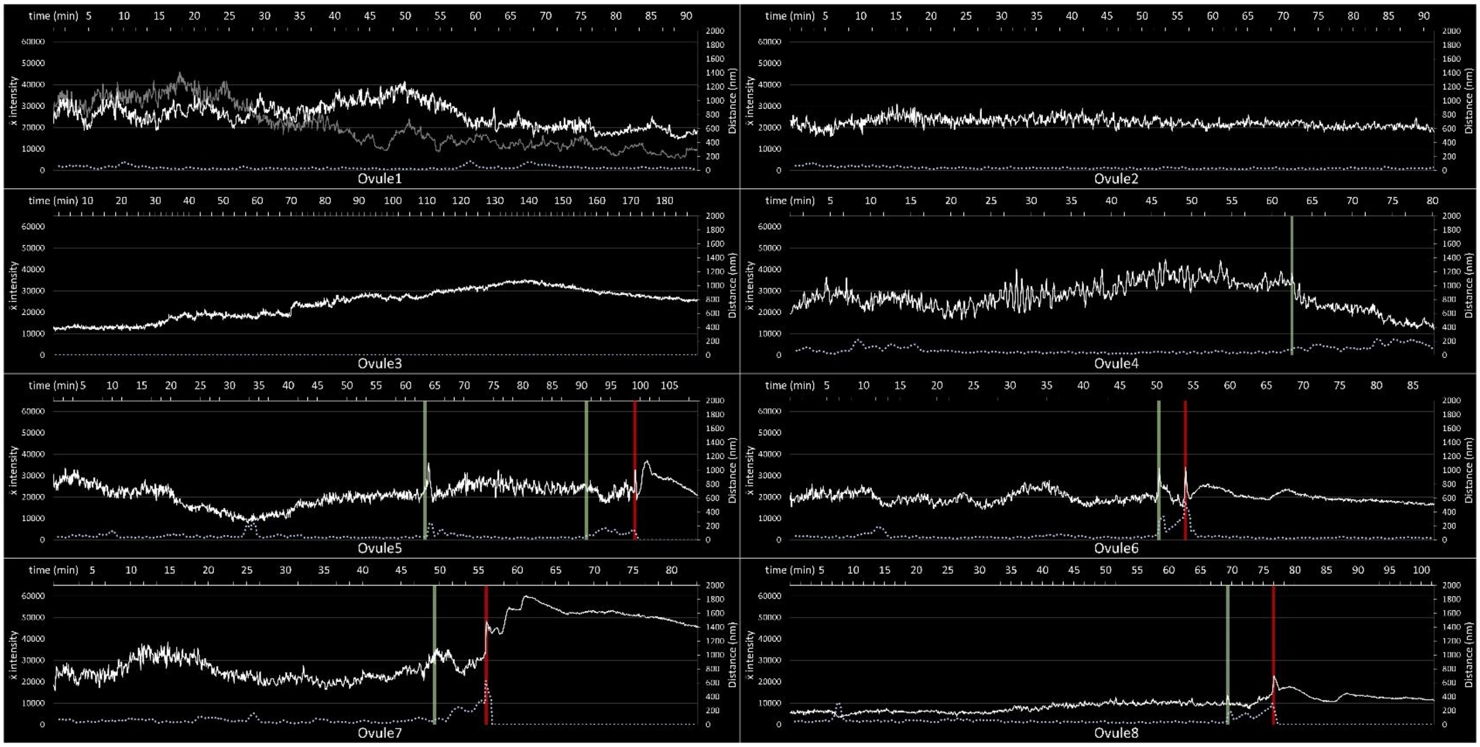


**Supplemental Figure 6.** ***aca9*** **PT R-GECO1 $[Ca^{2+}]_{cyt}$ traces during PT reception.**

Eight $[Ca^{2+}]$ traces of manual tip tracked R-GECO1-expressing *aca9* PTs interacting with *pMYB98:MARIS-GFP* synergids imaged every 5 seconds. The left Y axis is the mean intensity of summed projections of 16bit Z stacks and applies to the white line. The right Y axis is distance in nanometers and applies to the blue dotted line, which shows the moving average trendline (period 10) of the PT distance traveled calculated from the coordinates of the tip center of mass. The top X axis is time in minutes. The green line denotes the approximate transition point of the PT from phase I to phase II growth and the red line denotes rupture of the PT.

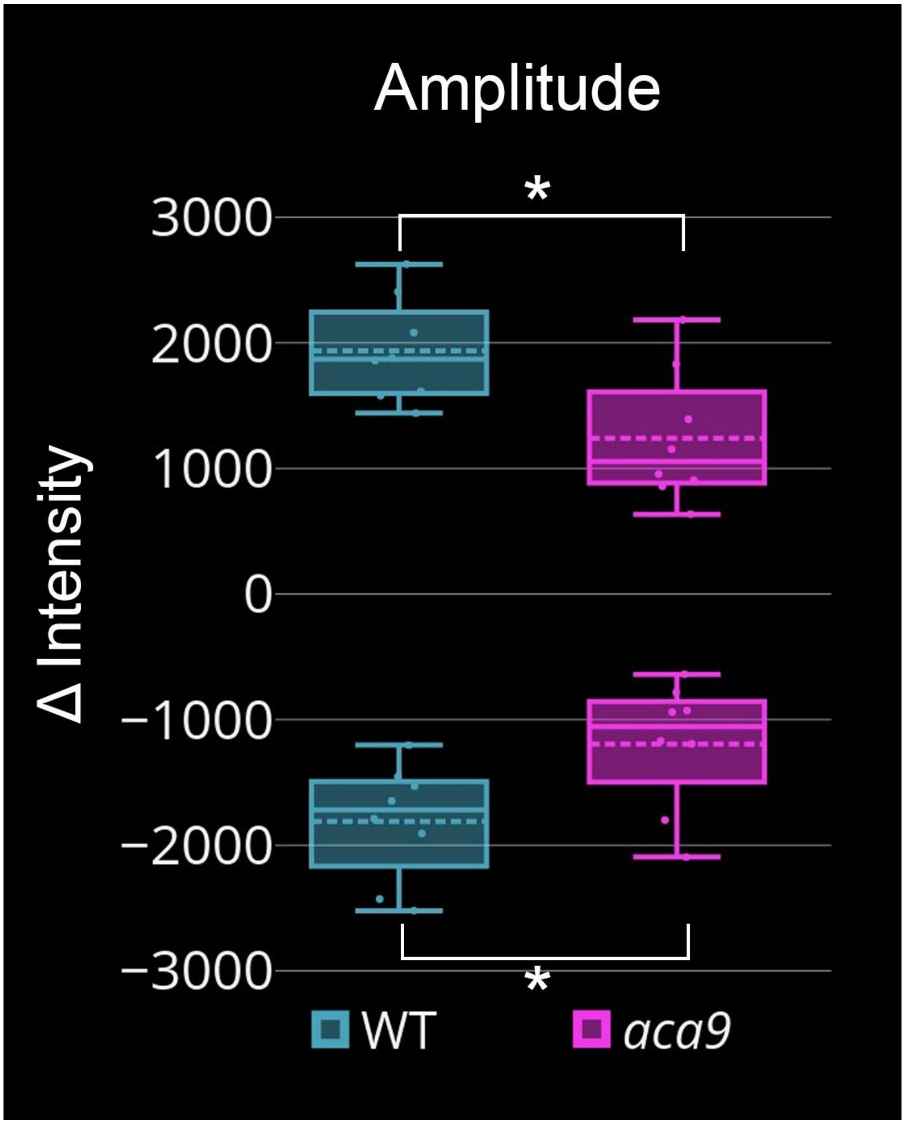


**Supplemental Figure 7. Amplitude of Phase I PT [$Ca^{2+}$]$_{cyt}$ spikes in WT and *aca9* during PT reception.**

Box plot of average Phase I [$Ca^{2+}$]$_{cyt}$ spike amplitude of eight WT *pLAT52:R-GECO1* (cyan) and *aca9 pLAT52:R-GECO1* PTs interacting with *pMYB98:MARIS-GFP* synergids during Phase I of PT reception (pink). The mean maxima and minima amplitude for WT PTs was 1936 AU and -1809 AU respectively, which was significantly different from *aca9* mean maxima, 1239 AU (p-value = 0.011) and minima, -1193 AU (p-value = 0.023) using a t-test.

**Supplemental Movie 1. PT tracking with widefield fluorescence microscopy.**

(A - L) Live imaging of PT reception showing manual tip tracking of PTs during guidance (white circles), phase I (orange circles), and phase II (green circles) aligned around the time of PT rupture. Yellow lines are outlines of thresholded synergid region of interests. Left of each panel shows just the stacked tracking outlines with time stamp (Hour: Min: Sec) and right shows just the RFP channel overlayed with synergid outline and PT tip tracking. Mean speed of the 9 tracked PTs (A-L) during guidance, phase I, and phase II were 1.36 µm/min, 0.47 µm/min, and 1.67 µm/min respectively. (J -L) show aberrant rupture of the synergids triggered during phase II of PT growth. Signal from the synergid cytoplasm was seen to be released into the central cell. Rupture point of the synergid is indicated with a red arrowhead. Video is looped 4 times for ease of viewing. Scale bars are 25 µm.

**Supplemental Movie 2. 3D reconstruction of *fer-1* synergid invaded by a PT.**

Video shows a 3D Z-projection of a *pFER:MARIS-GFP, fer-1* ovule (cyan) invaded by an *in-vitro* grown *pLAT52:dsRED* PT (orange). Video also show the segmentation and surface reconstruction of the synergid (pink) and PT (blue). Video is looped 2 times for ease of viewing. Time stamp is hr:min.

**Supplemental Movie 3. Live 3D imaging of peritubular membrane formation.**

(A) 3D model of ovule and segmented synergids of *pFER:FER-GFP* showing orientation of videos B and C that were imaged with X and Y as the ovules distal-proximal plane. (B, C) Live 3D imaging of *fer-1, pFER:MARIS-GFP* synergids (B) and *pMYB98:MARIS-GFP* synergids (C) interacting with a dsRED-expressing PT using SIV *cum septum*. Right sides show snapshots of the synergids at timepoint 0:00 and peritubular membrane formation after 24 minutes. (D) 3D model of ovule and segmented synergids of *pFER:FER-GFP* showing orientation of videos E and F that were imaged with X and Y as the ovules dorsal-ventral plane. (E, F) Live 3D imaging of *fer-1, pFER:MARIS-GFP* synergids (E) and *pMYB98:MARIS-GFP* synergids (F) interacting with a dsRED-expressing PT using SIV *cum septum*. Right sides show snapshots of the FA at timepoint 0:00 and peritubular membrane formation after 32 minutes. Scale bars are 10 µm. Videos are reversed and looped 6x for ease of viewing peritubular membrane formation. Levels of videos were adjusted and a gaussian blur applied to the GFP channel and smooth filter to the RFP channel.

**Supplemental Movie 4. Live 3D imaging of Synergid Actin and PT [$Ca^{2+}$] dynamics during cell invasion.**

(A, B, D, E) Live imaging of *pLAT52:R-GECO1* (Magenta) PT and *pFG:pLifeAct-HyPer7, pFER:FER-GFP* (Cyan) ovules interacting during PT reception using SIV *cum septum*. Merged channels (A, D) and just the GFP channels (B, E) are shown. (C, E) Animated graphs of the R-GECO1 signal intensity (white line) and distance of the PT (blue dotted line) corresponding to video A (C) and video D (F). Green bar denotes phase II, and red bar denotes PT rupture. Levels of the video were adjusted and a smooth filter applied to the RFP channel, but original images were unaltered for quantification. "Lysis" denotes rupture of the PT. Scale bars are 25 µm. Time stamp is hr:min:sec.

**Supplemental Movie 5. Live 3D imaging of PT [$Ca^{2+}$] dynamics during peritubular membrane formation.**

(A, B) Live imaging of *pLAT52:R-GECO1* (Magenta) PT and *pMYB98:MARIS-GFP* (Cyan) Synergids interacting during PT reception using SIV *cum septum*. Merged channels with region of interest used for intensity measurements (A) and just the GFP channel (B) are shown. An animated graph of the R-GECO1 signal intensity (white line) and distance of the PT (blue dotted line) is shown below. Orange bar denotes phase I, Green bar denotes phase II, and red bar denotes PT rupture. Levels of the video were adjusted and a gaussian blur applied to the GFP channel and smooth filter to the RFP channel, but original images were unaltered for quantification. "Lysis" denotes rupture of the PT. Scale bar is 10 µm. Time stamp is hr:min:sec.

**Supplemental Movie 6. Replicates of PT [$Ca^{2+}$] dynamics during peritubular membrane formation.**

Video shows eight replicates of R-GECO1-expressing PTs interacting with MARIS-GFP-expressing synergids using SIV *cum septum*. Each video is aligned 100 frames surrounding the phase transition spike. Videos also show the kymograph line used to generate the kymographs shown below each video which display just R-GECO1 (left) and overlay with MARIS-GFP (right). Scale bars are 10 µm. Levels of videos were adjusted and a gaussian blur applied to the GFP channel and smooth filter to the RFP channel, but were unaltered for quantification.

**Supplemental Movie 7. Live 3D imaging of PT [$Ca^{2+}$] dynamics during *fer-1* peritubular membrane formation.**

(A, B) Live imaging of *pLAT52:R-GECO1* (Magenta) PT and *fer-1, pMYB98:ROP2-GFP* (Cyan) synergids interacting during PT reception using SIV *cum septum*. Merged channels with region of interest used for intensity measurements (A) and just the GFP channel (B) are shown. An animated graph of the R-GECO1 signal intensity (white line) and distance of the PT (blue dotted line) is shown below. Levels of the video were adjusted and a gaussian blur applied to the GFP channel and smooth filter to the RFP channel, but original images were unaltered for quantification. Time stamp is hr:min:sec. Scale bar is 10 µm.

**Supplemental Movie 8. Replicates of PT [$Ca^{2+}$] dynamics during *fer-1* peritubular membrane formation.**

Video shows eight replicates of R-GECO1-expressing PTs interacting with MARIS-GFP-expressing (Ovules 1- 4) and ROP2-GFP-expressing (Ovules 5- 8) *fer-1* synergids using SIV *cum septum*. Each video is aligned 100 frames surrounding the phase transition spike. Videos also show the kymograph line used to generate the below kymographs which display just R-GECO1 (left) and overlay with MARIS-GFP or ROP2-GFP (right). Scale bars are 10 µm. Levels of videos were adjusted and a gaussian blur applied to the GFP channel and smooth filter to the RFP channel, but were unaltered for quantification.

**Supplemental Movie 9. Live 3D imaging of Synergid [$Ca^{2+}$] dynamics during PT reception.**

(A, B) Live imaging of *pLAT52:R-GECO1* (Magenta) PT and *pMYB98:GCaMP6f* (Cyan) Synergids interacting during PT reception using SIV *cum septum*. Merged channels with tip tracking (yellow circle) of the PT (A) and just the GFP channel (B) are shown. An animated graph of the change in signal intensity of the receptive synergid (white line, receptive synergid) and persistent synergid (blue line, pSYN) is shown below. Orange bar denotes phase I, Green bar denotes phase II, and red bar denotes PT rupture. Levels of the video were adjusted and a gaussian blur applied to the GFP channel and smooth filter to the RFP channel, but original images were unaltered for quantification. “Lysis” denotes rupture of the PT. Time stamp is hr:min:sec. Scale bar is 10 µm.

**Supplemental Movie 10. Live 3D imaging of *aca9* PT [$Ca^{2+}$] dynamics during PT reception.**

Videos of blocked (A, B) and WT-like (C, D) *aca9, pLAT52:R-GECO1* (Magenta) PTs and *pMYB98:MARIS-GFP* (Cyan) synergids interacting during PT reception using SIV *cum septum*. Merged channels with region of interest used for intensity measurements (A, C) and just the GFP channels (B, D) are shown. Levels of the video were adjusted and a gaussian blur applied to the GFP channel and smooth filter to the RFP channel, but original images were unaltered for quantification. “Lysis” denotes rupture of the PT. Time stamps are hr:min:sec. Scale bars are 10 µm.

**Supplemental Movie 11. Replicates of *aca9* PT [$Ca^{2+}$] dynamics during peritubular membrane formation.**

Video shows eight replicates of R-GECO1-expressing *aca9* PTs interacting with MARIS-GFP-expressing synergids using SIV *cum septum*. Each video is aligned 100 frames surrounding the phase transition spike (no transition in ovules 1-3). Videos also show the kymograph line used to generate the kymographs below each video that display just R-GECO1 (left) and overlay with MARIS-GFP (right). Scale bars are 10 µm. Levels of videos were adjusted and a gaussian blur applied to the GFP channel and smooth filter to the RFP channel, but were unaltered for quantification.

**Supplemental Movie 12. Live 3D imaging of Synergid and *aca9* PT [$Ca^{2+}$] dynamics during PT reception.**

(A, B) Live imaging of *aca9 pLAT52:R-GECO1* (Magenta) PT and *pMYB98:GCaMP6f* (Cyan) Synergids interacting during PT reception using SIV *cum septum*. Merged channels with tip tracking (yellow circle) of the PT (A) and just the GFP channel (B) are shown. Below shows an animated graph of the change in signal intensity of the right synergid (blue line), left synergid (white line, pSYN), and PT (purple line). Orange bar denotes onset of phase I. Levels of the video were adjusted and a gaussian blur applied to the GFP channel and smooth filter to the RFP channel, but original images were unaltered for quantification. Scale bar is 10 µm. Time stamp is hr:min:sec.
